# The Impact of Noise on Memorability in Minds and Machines

**DOI:** 10.64898/2026.10.19.759153

**Authors:** Huiyu Ding, Martin Wiener

## Abstract

Some images are consistently remembered better than others, a stimulus property known as memorability, and deep neural networks (DNNs) are increasingly used to estimate it in place of human observers. A recent account proposes that memorable images are processed more efficiently. We argue that memorability can also be viewed as signal quality, and that efficient processing should therefore be robust to noise. Comparing memorability estimates from state-of-the-art (SOTA) models with human performance in continuous recognition tasks, we found that the two consistently diverged in how memorability changed as images were degraded by increasing levels of Gaussian noise and blur, such that models under-estimated high memorability images with noise, while over-estimating low memorability images with noise, a pattern resembling stochastic resonance. In contrast, noise degraded memorability estimates in humans, yet high memorability images were resistant to this effect. We discovered this divergence arose because conventional models are influenced by how noise alters their attention across images, whereas humans learn how images relate to one another. Building on these findings, we developed GraphMem, a model that estimates memorability from the relational structure among images rather than from the features of individual images. GraphMem outperformed SOTA models in estimating memorability under noise and blur and aligned most closely with human data. These findings caution against relying on conventional DNN-based memorability estimates for novel images, and suggest that modeling the relational structure in vision models yields more human-like memorability estimates.

## Introduction

Glance at a photograph of a child blowing out birthday candles and one of an empty parking lot, and a day later you will likely remember the image of a birthday party but not the parking lot. Strikingly, most people tend to remember or forget the same images, an effect known as image memorability (Bainbridge et al., 2013; Isola et al., 2011). Yet, we rarely see visual information under ideal conditions; we glimpse scenes through a rain-streaked windshield, on grainy security footage, or in the dim light of evening. Would the birthday photograph still be remembered better if it were seen through visual noise? Could noise make an image more, or less, memorable? In this study, we investigate how visual noise changes image memorability in state-of-the-art (SOTA) deep neural network models (DNNs) and human observers and seek to develop a new model that outperforms existing models in noise tolerance.

To quantify how well an image is remembered later, memorability scores have commonly been estimated using a continuous recognition task, in which participants are presented with a sequence of novel and repeated images and asked to identify the repeats (Bainbridge et al., 2013; Bylinskii et al., 2015; Isola et al., 2014; Khosla et al., 2015). The memorability measurement can be based on various memory metrics, such as hit rate (Bainbridge et al., 2013; Isola et al., 2011), sensitivity (*d′*; Bainbridge, 2017), or corrected recognition (hit rate minus false alarm rate; Bainbridge & Rissman, 2018). While human memory is multiply determined, it has been shown that image memorability is an intrinsic property of images. For example, people exhibit high consistency in what items they remember and forget for a wide range of visual stimuli, with split-half Spearman’s correlation across participant samples as high as *ρ* = .75 (for a review, see Bainbridge et al., 2026). Additionally, convolutional neural network (CNN) models trained on images and image labels produce memorability estimates that closely align with human performance (Khosla et al., 2015; Needell & Bainbridge, 2022). The factors that contribute to image memorability, however, remain debated. For example, image memorability cannot be reduced to individual image features, as differences in memorability persist when low-level features (e.g., color, spatial frequency) or high-level features (e.g., scene content, attractiveness) are controlled (Bainbridge et al., 2017; Bylinskii et al., 2015; Dubey et al., 2015; Isola et al., 2011, 2014). Image memorability is also not strongly correlated with subjective judgments of images, such as aesthetics or interestingness (Isola et al., 2014), and has been shown to be independent of top-down factors such as depth of encoding or cognitive control strategies (Bainbridge, 2020; Wakeland-Hart et al., 2022). Memorability is also separate from cognitive factors that typically modulate memory performance, such as attentional capture, as memorable images do not automatically capture attention in oddball visual search tasks (Bainbridge, 2020).

One prominent account of why some images are consistently more memorable than others proposes that memorable images are processed more efficiently (Bainbridge et al., 2026). On this view, memorable images are more readily perceived, categorized, and encoded because their visual and semantic information can be integrated rapidly and at relatively low cognitive cost. In particular, memorable images may have a “good” perceptual organization, in which their elements are quickly bound into a coherent representation and matched to existing templates, facilitating the extraction of meaning. Several lines of evidence support this account. At the perceptual level, memorable images reached detection threshold faster than forgettable images in an intact/scrambled detection task (Deng et al., 2024), and a comparable pattern has been reported when contrasting typical and atypical images (Center et al., 2022). Memorable images also yield above-chance recognition and retain an advantage over forgettable images at presentation durations as short as 13 ms (Broers et al., 2018; Ye et al., 2024). At the neural level, memorability can be decoded from magnetoencephalography signals shortly after stimulus onset (Lahner et al., 2024; Mohsenzadeh et al., 2019). Beyond initial perception, this processing advantage extends to visual working memory. Gillies and colleagues (2023) presented memorable and forgettable images in sets of varying sizes that were either mixed or pure (only memorable or only forgettable images). Pure sets of memorable images were better maintained in working memory than pure sets of forgettable images, suggesting that memorable images are represented more efficiently in capacity-limited working memory. In mixed sets, memorable images also attracted more working memory resources than the forgettable images they competed with. Finally, at the computational level, Ma and colleagues (2024) found that a recurrent convolutional network model (BLnet) converged on a classification of memorable images faster, with its classification uncertainty (entropy) declining more rapidly across recurrent time steps, suggesting that memorable images are processed faster. Taken together, converging evidence suggests that memorable images are processed more rapidly and efficiently than forgettable images.

Notably, nearly all this evidence comes from clean, high-quality images viewed under ideal conditions. If memorability reflects, at least in part, how efficiently an image’s signal is extracted and processed at the perceptual level, then memorability should also depend on the quality of that signal. From a signal-processing perspective, efficient processing depends not only on speed but also on robustness; that is, on how reliably relevant information can be extracted from the noise that accompanies it. Adding noise to visual stimuli offers a direct test of this robustness, and studies have shown that humans and non-human primates can detect and identify objects embedded in substantial levels of visual noise (Gold et al., 1999; Jang et al., 2021; Kwon & Legge, 2011; Pratte et al., 2013), whereas DNNs tend to fail catastrophically when humans do not (Dodge & Karam, 2017; Geirhos et al., 2018; Jang et al., 2021; Jang & Tong, 2021, 2024). This human–machine gap matters for memorability research. DNN-based memorability models are increasingly used in place of human raters to score, select, and even manipulate images (e.g., Goetschalckx et al., 2020; Khosla et al., 2013; Needell & Bainbridge, 2022; Younesi & Mohsenzadeh, 2026), yet their agreement with human memorability has been validated almost exclusively on clean images. If these models fail to accurately track memorability judged by humans, the validity of DNN-based estimates of human memorability needs to be reexamined, and comparing models and humans with the same noise-embedded images is an essential first step.

How then might noise affect memorability? Contrary to the common misconception that noise-embedded visual processing is suboptimal and problematic, studies showed that adding a moderate amount of noise can enhance the detection of weak and subthreshold signals, a phenomenon known as stochastic resonance (Collins et al., 1996; Faisal et al., 2008; McDonnell & Abbott, 2009; Moss et al., 2004; for a review, see Ward et al., 2002). Critically, stochastic resonance predicts that the effect of noise depends on the initial strength of the signal. Signals that are already strong are degraded by added noise, whereas weak signals may be pushed above threshold and thereby benefit from it. Applied to memorability, this raises the possibility that the effect of noise may behave similarly. Specifically, adding noise may lower the memorability of initially highly memorable images, whereas initially less memorable images may benefit from it. It is important to note, however, that some studies have shown that noise degradation affects perception and memory differently. For example, Meng and Potter (2008) presented observers with rapid sequences of partially occluded scenes and found that occluding up to 30% of each image left target perception largely intact but significantly impaired subsequent recognition memory. It therefore remains an empirical question whether, and how, memorability varies with added noise.

To explore how memorability varies in the face of noise, we first used DNNs known to predict human memorability to estimate the memorability of naturalistic scenes perturbed with increasing levels of Gaussian noise. Second, we compared these estimates with human recognition memory for the same images in a continuous recognition task. To preview our results, the DNNs showed a pattern resembling stochastic resonance where noise lowered the estimated memorability of initially highly memorable images but raised that of initially less memorable images. Human observers, however, showed a divergent pattern. Recognition of highly memorable images was robust to noise, whereas recognition of less memorable images declined. This divergence persisted across additional SOTA models, including models trained on noise-perturbed images. Because image typicality is related to memorability (Kramer et al., 2023), we also examined typicality and found that noise altered how similar images were to one another, suggesting the importance of considering local context. We therefore developed *GraphMem*, a graph-based model that estimates memorability from an image’s similarity to other images in the dataset. GraphMem outperformed all models in their ability to match human performance, and this advantage extended to images degraded by blur.

## Results

### Effects of noise on model-estimated memorability

We begin addressing the question of how conventional memorability models estimate image memorability under noise by using ResMem (Needell & Bainbridge, 2022) on the LaMem dataset, for 196 scenes with uniform sampling of memorability separated into seven bins, each presented at five levels of Gaussian noise (980 images total; **Fig. 1**a, d; Methods). Fitting slopes of predicted memorability across five noise levels with a one-way ANOVA, we found a significant main effect of bin, *F* (6,189) = 18.68, *p* < .001, η^2^_p_ = 0.37. To characterize the direction of this effect, by-image slope values were fit to a linear regression model with numerical memorability bin as a fixed effect, which yielded a significant fixed effect, *β* = 0.12*, SE* = 0.03*, t* = 4.65*, p* < .001, with high memorability images characterized by a negative slope and high memorability images characterized by a positive slope, such that low-memorability images were estimated as more memorable with noise, while high memorability images were estimated as less memorable, resembling the stochastic resonance pattern.

To determine if this pattern was due to the specific model structure, we tested the same images through two additional memorability models: ViTMem (Hagen & Espeseth, 2023) and AMNet (Fajtl et al., 2018). Both models reproduced the effect (**ViTMem:** **Fig. 1c**, *F* (6,189) = 7.43, *p* < .001, η^2^_p_ = 0.19; AMNet: **Fig. 1b**, *F* (6,189) = 36.74, *p* < .001, η^2^_p_ = 0.54), where predicted memorability decreased with noise for high-memorability images and increased for low-memorability images. The convergence of results across transformer-based and convolutional architectures suggests that the effect is not specific to a particular model class and instead reflects a systematic relationship between image noise and the visual features these models use to predict memorability.

Next, we investigated why noise affected memorability differently across images by examining attention maps from AMNet (Fajtl et al., 2018). The dispersion of attention the model used in estimating memorability was operationalized by entropy values of attention maps at the final LSTM step (Methods; **Fig. 1e–g**). A one-way ANOVA on entropy slopes yielded a significant main effect, *F* (6, 189) = 4.74, *p* < .001, η^2^_p_ = 0.15. A linear regression model on entropy slopes yielded a significant fixed effect of bin, *β* = -0.2*, SE* = 0.01*, t* = -4.16*, p* < .001, such that high-memorability images were typically characterized by a central focal point driving memorability, but with noise became dispersed throughout the image thus lowering memorability. In contrast, the dispersion remained relatively stable with noise for low-memorability images (bin 5: b = 0.02, *p* = 1; bin 6: *b* = −0.05, *p* = 1; bin 7: *b* = −0.08, *p* = .775). Thus, attentional dispersion can partially explain the stochastic resonance pattern observed with DNNs and why noise lowered the estimated memorability of highly memorable images.

**Fig. 1.**
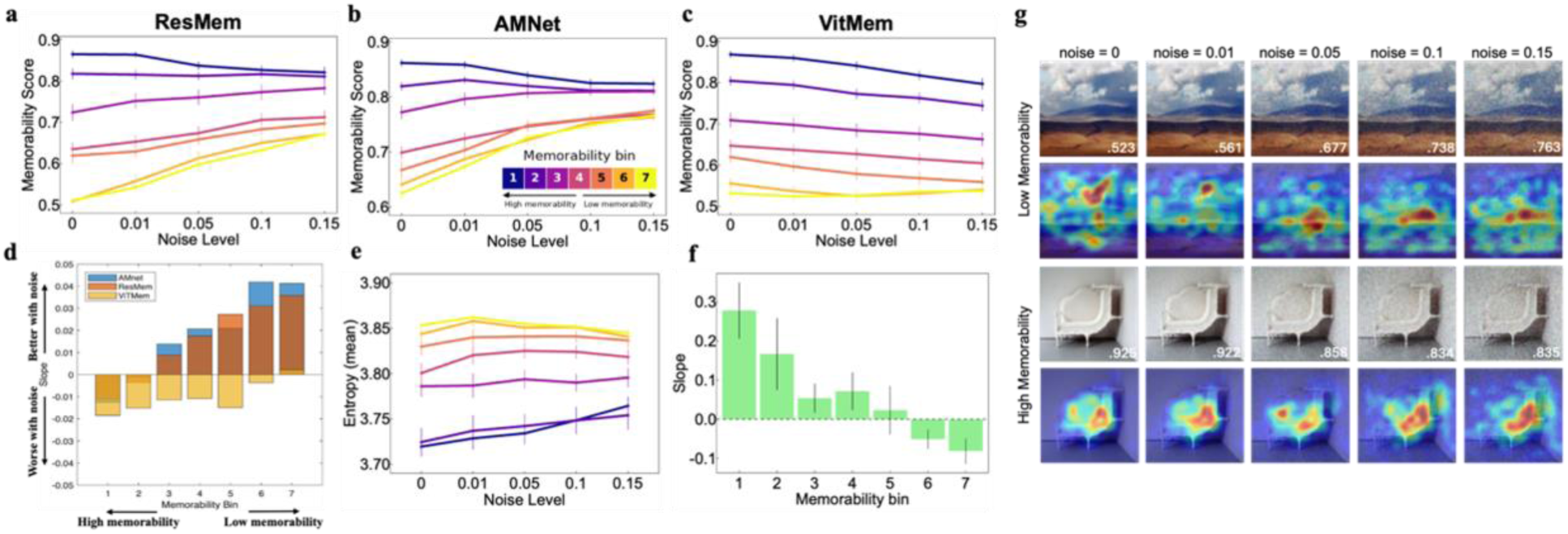
Effects of noise on model-estimated memorability. Estimated memorability with varied noise levels from ResMem (**a**), AMNet (b), and ViTMem (**c**). Model-predicted memorability decreased with increased noise levels for high-memorability images but increased for low-memorability images (**d**). **e**, **f**, Entropy of AMNet attention maps increased with noise for high-memorability images, whose attention was initially concentrated on a central focal point. **g**, Example AMNet attention maps at each noise level for two images; the top row displays a low memorability image, characterized by a distributed attention map, that with noise becomes more concentrated in a single location and thus raises the memorability score, whereas the bottom row displays a high memorability image, characterized by a focused attention map, that with noise becomes more distributed across the image and thus lowers the memorability score. Error bars denote SEM.

### Effects of noise on human recognition memory

Our results so far demonstrate that DNNs reliably show an interaction between memorability and noise added to images. As noise increases, estimated memorability for initially highly memorable images decreases, whereas estimated memorability of initially less memorable images benefits from noise degradation. Because these models were trained to approximate human memorability, it remains an open question whether human observers show the same pattern. If noise reshapes human encoding the way it redistributes the models’ attention (as seen with the AMnet attention pattern), noise degradation should selectively benefit initially weak images, and recognition slopes should vary systematically with memorability bins. To test this, we ran the same image set in a continuous recognition experiment with human participants.

To explore whether human participants exhibit a stochastic resonance effect, corrected recognition (CR) rate was calculated for each participant by subtracting the false alarm rate from the hit rate for target images. Behavioral results across memorability bins and noise levels, together with the slope values, are visualized in **Fig. 2**b–e. Slopes of hit rate (HR), false alarm rate (FA), and CR across noise levels were analyzed for each participant. Hit rates and false alarm rates were negatively correlated (*r* = −0.87). The one-way repeated-measures ANOVA on by-subject slope values across noise levels yielded a significant main effect of bin for FA (**Fig. 2c**), *F* (6,522) = 4.60, *p* < .001, η^2^_p_ = 0.01, suggesting that the likelihood of making incorrect responses to low-memorability images increased with greater levels of noise to a greater degree than high-memorability images, *β* = 0.12*, SE* = 0.03*, t* = 4.65*, p* < .001. We did not find a significant main effect of bin for HR (**Fig. 2b**), *F* (6,522) = 1.18, *p* = .315, but there was a significant main effect of bin for CR (**Fig. 2d, e**), *F* (6,522) = 2.96, *p* = .007, η^2^_p_ = 0.07. Linear mixed effect model on CR yielded a significant fixed effect of bin, *β* = -0.17*, SE* = 0.05*, t* = 3.36*, p* < .001, suggesting that recognition of more memorable images was more resistant to increasing noise with flatter slopes whereas less memorable images were more negatively affected by image degradation Crucially, this pattern diverged from estimated memorability generated by conventional models.

**Fig. 2.**
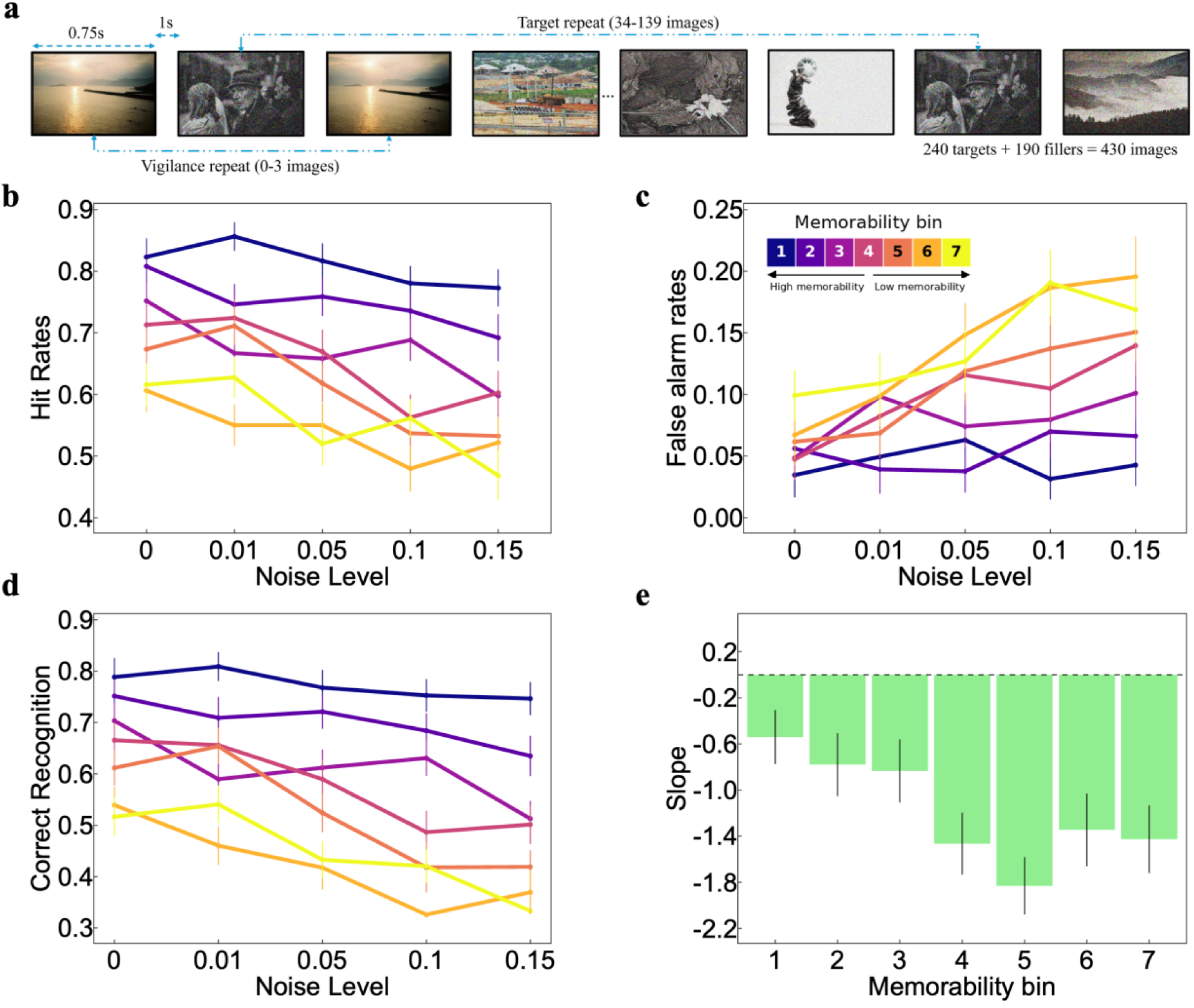
Effect of noise on human recognition memory. **a**, Participants (n = 88) completed a continuous recognition memory task with stimuli drawn from the LaMem dataset with different levels of noise perturbation, with each subject receiving each image at only one level of perturbation. We found a significant main effect of memorability bins for false alarm rate (**c**) but not for hit rates (**b**). Corrected recognition remained stable across increasing noise for high-memorability images but declined for low-memorability images (**d**). **e**, Subject-level slopes of corrected recognition across noise, plotted by memorability bin. Error bars denote SEM.

### The human–model divergence persists across additional vision models

The divergence between model estimates and human performance could reflect our particular choice of models rather than a general property of vision models. That is, while the three models we employed here have been trained for memorability, they rely on architectures that have since given way to newer ones. Therefore, we trained several additional state-of-the-art (SOTA) vision models on the LaMem dataset and compared their estimates with the human data (referred to as “NoiseMem”). Additionally, prior studies showed that re-training vision models with noise-treated stimulus sets (e.g., blur and Gaussian noise) could improve models’ ability to recognize degraded images (Vasiljevic et al., 2016; Zhou et al., 2017) and to predict human behavioral performance, as well as increase their alignment with the human visual system (e.g., Jang et al., 2021; Jang & Tong, 2024, 2025; Thompson, 2021). We therefore retrained all SOTA models, together with the three models used previously (ResMem, ViTMem, and AMNet), on images randomly perturbed with Gaussian noise (0–0.15), reasoning that exposure to noise might improve memorability prediction under noise degradation. Results on Spearman correlation from all models, trained with or without noise, to LaMem estimated memorability and human performance (NoiseMem) are reported in **Table 1**. None of the models could capture the pattern found in human performance, with all exhibiting a stochastic resonance effect to varying degrees. Furthermore, training with noise-perturbed images did not significantly improve the models’ ability to estimate LaMem memorability, and some models even performed worse (e.g., ResMem). Likewise, training with noise did not improve the models’ prediction of human recognition performance; in fact, with few exceptions (VitMem, ConvNeXt), most models performed worse overall with noise training. When model-estimated memorability was correlated with mean human CR rates, all models performed poorly in capturing the general direction of the human performance pattern (see Slope Effect in Table 1). In contrast, GraphMem, a context-sensitive model we introduce below, showed the best alignment with human performance.

**Table 1.** Spearman correlations between model-estimated memorability and both LaMem ground-truth memorability and human recognition performance (NoiseMem), for each vision model trained on clean images and on noise-perturbed images (Gaussian noise, 0–0.15). ρ = Spearman rank correlation. *MSE* = mean squared error. The NoiseMem effect correlates the average 5x7 matrices of memorability scores between model output and human performance. Slope Effect is the Spearman correlation, across seven memorability bins, between the noise slopes of model-estimated memorability and those of human corrected recognition, where a noise slope is the change in score with increasing noise level within a bin. Positive values indicate that the model reproduces the directional human pattern of noise effects across memorability levels, whereas more negative values indicate the model produced a stochastic resonance-like effect. The Slope Effect for noise-trained GraphMem (ρ = 0.57, 95% bootstrap CI [0.29, 0.75]) was calculated by averaging the scores following 100 permuted training runs with random assignment of noise level to images (mirroring 100 human participants).

| Trained without noise |  | Trained with noise |  | NoiseMem |  | SlopeEffect |  |  |
| --- | --- | --- | --- | --- | --- | --- | --- | --- |
| Model | $\rho$ | $MSE$ | $\rho$ | $MSE$ | Without noise | With noise | Without noise | With noise |
| Vit_DINOv3 | .68 | 0.01 | .16 | 0.01 | 0.77 | 0.31 | −0.68 | −0.54 |
| VitMem | .69 | 0.01 | **.65** | 0.01 | 0.82 | 0.83 | −0.75 | −0.75 |
| ResMem | .64 | 0.01 | −.01 | 0.01 | 0.82 | 0.21 | −0.68 | −0.57 |
| Eva02 | .67 | 0.01 | .61 | 0.01 | 0.65 | 0.57 | −0.82 | −0.68 |
| Vit_MAE | .64 | 0.01 | .61 | 0.01 | 0.73 | 0.71 | −0.75 | −0.68 |
| Vit_CLIP | .67 | 0.01 | .42 | 0.01 | 0.75 | 0.64 | −0.68 | −0.75 |
| ConvNeXt | **.69** | 0.01 | .63 | 0.01 | 0.77 | 0.82 | −0.75 | −0.75 |
| AMNet | .65 | 0.01 | .63 | 0.01 | 0.59 | 0.59 | −0.75 | −0.75 |
| GraphMem | .48 | 0.01 | .36 | 0.02 | **0.85** | **0.84** | **−0.11** | **0.57** |

### Noise perturbation modulates image typicality

One possible reason that the aforementioned SOTA models diverged from human performance could be that these models were all trained and tested on images in isolation. Yet, image memorability likely depends not just on a given image, but on the larger context in which that image was presented. Indeed, recent research has noted that estimated memorability may be inverted when the composition of the training dataset is manipulated, suggesting a context-dependent memorability effect (Nosofsky et al., 2025; Simpson et al., 2026). It is thus possible that images with varying levels of memorability were differentially changed relative to one another with increasing noise. In support of this possibility, we measured the pixel-level typicality of our images at each noise level. We found that memorability inversely correlated with typicality for clear images (*r* = -0.18, *p* < .001), with more memorable image being less typical. Importantly, we found that noise generally decreased image typicality scores, but the effect varied as a function of initial memorability score (**Fig. 3a, b**), *F* (6,189) = 2.94, *p* = .009, η^2^_p_ = 0.01. Specifically, high-memorability images (bin 1: *b* = −0.43; bin 2: *b* = −0.47; bin 3: *b = −0.46) were more resistant* to this effect than low memorability images (bin 4: b = −0.45; bin 5: *b* = −0.65; bin 6: *b* = −0.80; bin 7: *b* = −0.70), *β* = -0.06, *SE* = 0.02, *t* = 3.86, *p* < .001. As a result, any model of memorability, in the face of noise, needs to account for inter-image effects to match human performance.

To address this issue, we developed a new model, called “GraphMem” (**Fig. 3c**), which predicts an image’s memorability from its neighborhood in a similarity graph built over all LaMem images (for more details, see Methods). We found estimated memorability from GraphMem was largely resistant to noise (**Fig. 3d**) and did not vary as a function of memorability bins, *F* (6,189) = 1.30, *p* = .259, η^2^_p_ = 0.04, mirroring human data (**Fig. 3e**). Following similar procedures in relating conventional models with NoiseMem, GraphMem achieved the best alignment in general, whether trained with or without noise, demonstrated by the highest Spearman correlation of subject-level slope averages across memorability bins and noise levels (Table 1, NoiseMem column). GraphMem also came closest to reproducing the direction of human noise effects across memorability bins. When trained on clean images, its Slope Effect was the least negative of all models (ρ = −0.11), and after fine-tuning with noise it was the only model with a positive Slope Effect (ρ = 0.57, 95% bootstrap CI [0.29, 0.75]; **Table 1**). However, GraphMem was overall worse at predicting individual memorability measures than conventional models, an important future direction that we revisit in the Discussion.

**Fig. 3.**
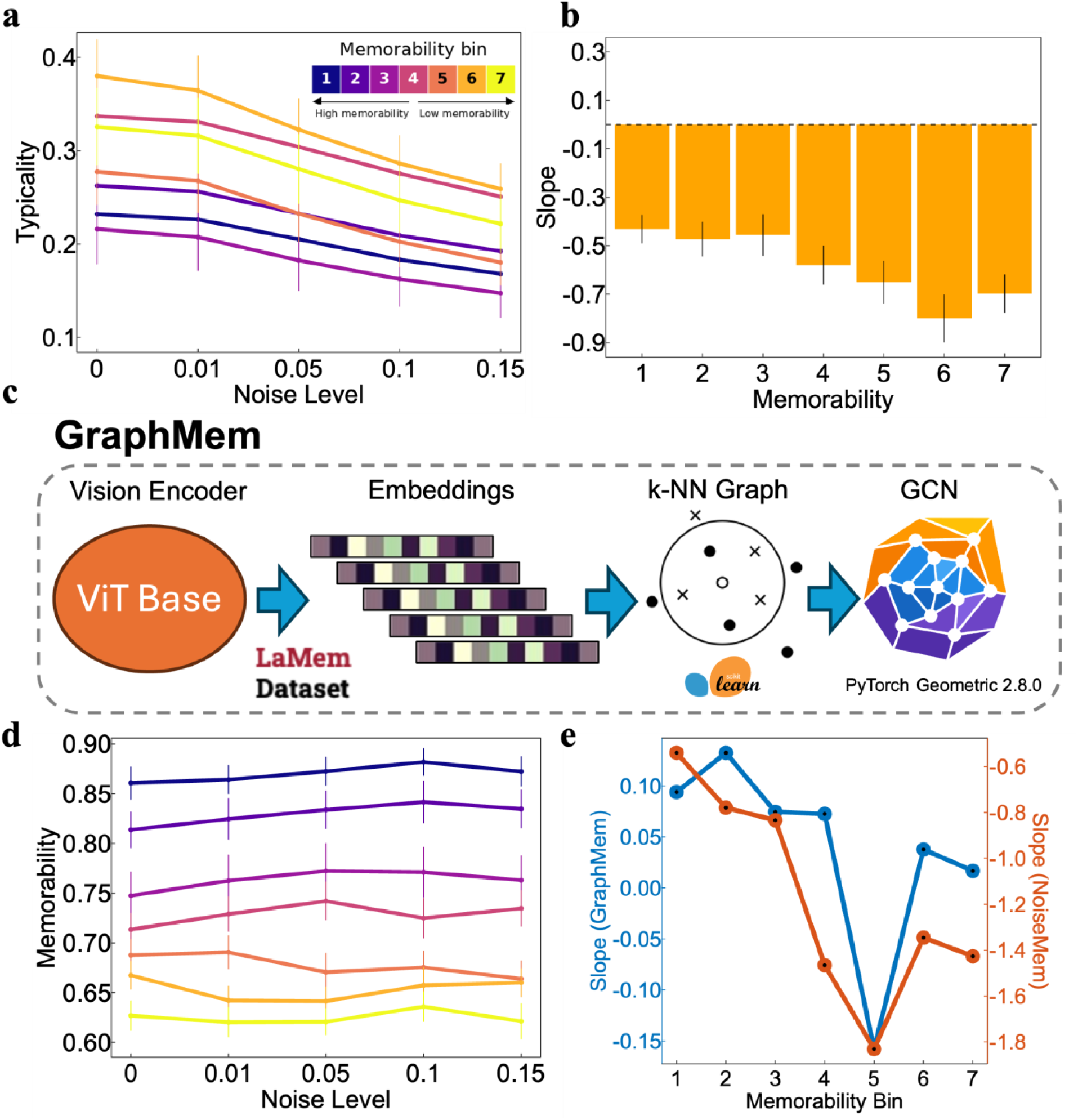
Effect of noise on image typicality and GraphMem. **a,** Image typicality as a function of noise level, by memorability bin. **b**, Slope estimate of typicality across noise, plotted against memorability. **c**, GraphMem architecture. **d**, GraphMem-estimated memorability across noise levels, by memorability bin. **e**, Slopes of estimated memorability across noise by bin for GraphMem (blue, left axis) and for the human data (NoiseMem; orange, right axis). Error bars denote SEM.

### GraphMem matches human performance with blurred images

We have so far established that GraphMem achieved the best performance in matching human data in terms of predicting memorability with noise-added images by taking into account the contextual effect. Next, we aimed to extend the results to visual blur. Blurry images are commonly encountered in everyday human vision (Burge & Geisler, 2011; Sprague et al., 2016), and studies showed that, similar to Gaussian noise applied to images, CNNs tend to fail on blurry images where humans do not (e.g., Dodge & Karam, 2017; Geirhos et al., 2018). Therefore, it is of theoretical and practical importance to apply our methods to images perturbed with blur to explore whether we would observe similar patterns. To that end, we conducted a second online experiment (referred to as BlurMem) that followed the procedure of NoiseMem but perturbed images with varying degrees of blur. We then compared human performance with memorability estimated from ResMem and noise-trained GraphMem. This allowed us to test whether GraphMem could outperform established vision models with a different type of image degradation and to assess the generalizability of training on one type of degradation (Gaussian noise) to another (blur).

Participants completed a continuous recognition task with images degraded with different levels of blur (image examples in **Fig. 4a**). Hit and FA rates were negatively correlated (*r* = -0.77) A one-way repeated-measures ANOVA on HR (**Fig. 4b**) yielded a significant main effect of bin, *F* (6,540) *=* 3.30*, p =* .003, η^2^_p_ = 0.04. A linear mixed effect model yielded a significant fixed effect of bin, *β* = 0.02, *SE* = 0.01, *t* = 3.80, *p* < .001, demonstrating that initially low-memorability images were more resistant to the effect of blur (than high-memorability images. Model fit to FA (**Fig. 4c**) also yielded a significant main effect of bin, *F* (6,540) = 2.25, *p* = .037, η^2^_p_ = 0.02, where participants made more incorrect responses as noise level increased for low-memorability images than high-memorability images, *β* = 0.01, *SE* = 0.01, *t* = 2.29, *p* =.022. Corrected recognition (**Fig. 4d**) was overall negatively affected by blur and showed an emerging significant main effect of bin, *F* (6,540) = 1.89, *p* = .081, η^2^_p_ = 0.02. This pattern differed from the effect of Gaussian noise on memorability, potentially due to the pervasive presence of blur in everyday visual processing (i.e., parafoveal defocused blur) and the fact that humans can leverage blur to support various visual recognition tasks (Bar, 2004; Held et al., 2012; Oliva & Torralba, 2007).

### GraphMem outperforms in estimating memorability with blurred images

To explore whether GraphMem outperforms other models with blurred images, we used GraphMem and ResMem to estimate the image memorability across levels of blur (**Fig. 4e–g**). We again found that ResMem diverged from the pattern observed with human data with a significant main effect, *F* (6,189) = 29.24, *p* < .001, η^2^_p_ = 0.48, where the estimated memorability of initially high-memorability images was negatively affected by blur perturbation but initially low-memorability images benefitted from blur, as confirmed by a significant positive effect of memorability bin on blur slopes, *β* = 0.01, *SE* = 0.01, *t* = 12.98, *p* < .001 GraphMem, on the other hand, did not show the stochastic resonance pattern but showed a general negative effect of blur on memorability, *F* (6,189) = 0.84, *p* = .540, η^2^_p_ = 0.03, more closely matching human data.

**Fig. 4.**
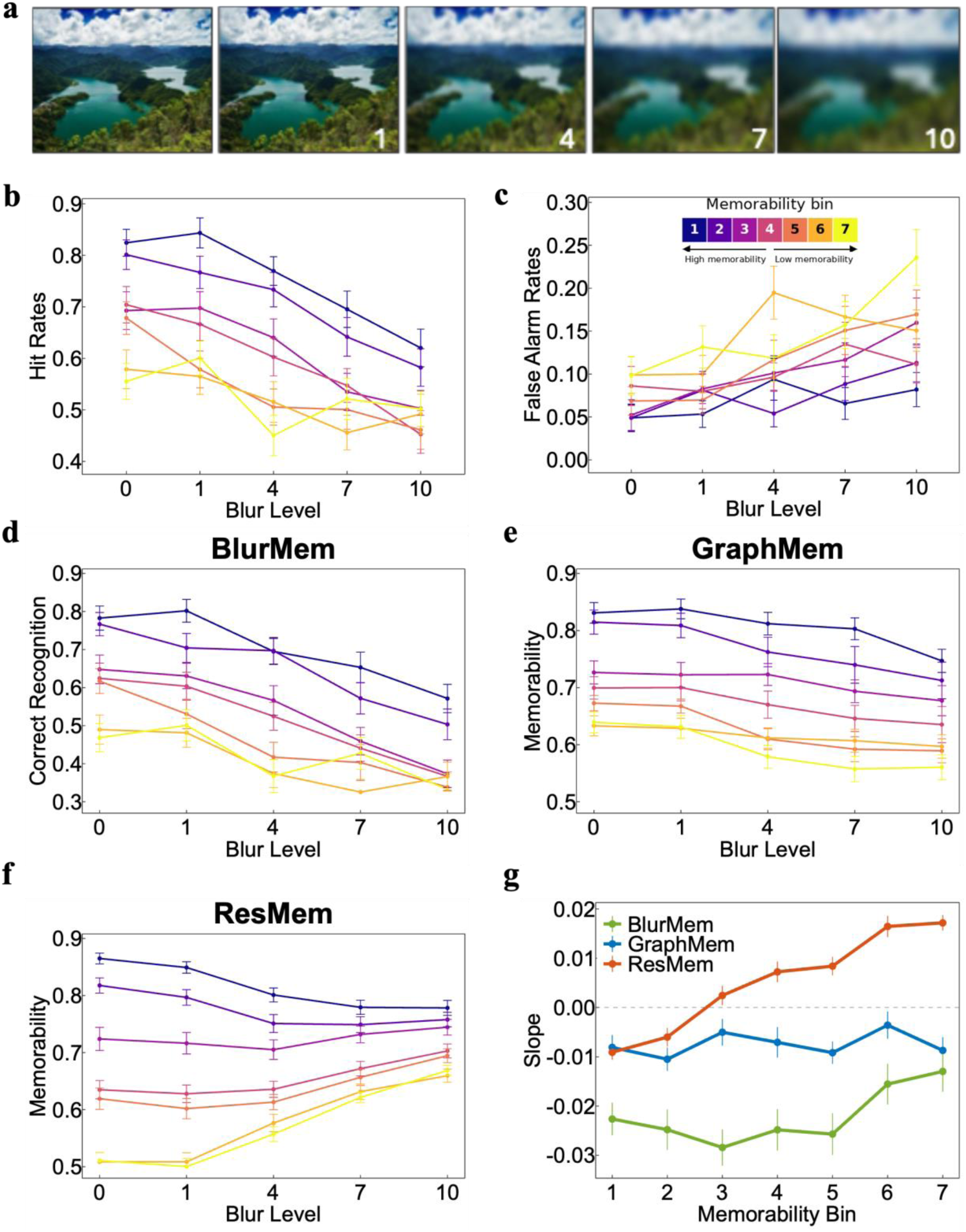
Human recognition and model estimation on blurred images. **a**, Example images at five blur levels (0–10). Hit rates (**b**), false alarm rates (**c**), and corrected recognition rate (**d**) across blur levels as a function of memorability bins. Estimated memorability for blurred images across memorability bins from noise-trained GraphMem (**e**) and ResMem (**f**). **g**, Slope of estimated memorability across blur levels for each memorability bin. GraphMem more closely matches the pattern observed with human participants than ResMem. Error bars denote SEM.

## Discussion

Memorability is argued to be an intrinsic property of images, and deep neural network (DNN) models have been developed to accurately estimate image memorability. Here, we evaluated how DNNs estimated image memorability with noise-perturbed images and how these models compare with human observers. We found that conventional state-of-the-art memorability models produced divergent patterns from human observers, observed for both Gaussian noise and blur. This human–model divergence was not resolved by fine-tuning models with noise-perturbed images. GraphMem, however, best matched the human pattern of noise effects. These findings suggest that accounting for the relational structure of the image set may help models better capture human memory under noise.

Conventional memorability models responded to noise in a consistent but non-human-like way. The existing SOTA models, regardless of the model structures and backbones, reliably captured initial memorability and exhibited a commonly observed stochastic resonance pattern (Stocks, 2000), where the direction of estimated memorability across noise levels varied as a function of initial memorability. Alternatively, this pattern could reflect simple regression to the mean: as noise removes diagnostic information, model estimates drift toward the average. However, our attention-map results reveal a shift in the image regions that models weight when estimating memorability, suggesting that noise perturbation changes how models process images rather than merely degrading their estimates. Moreover, when taking context into account, GraphMem did not exhibit the same pattern, further arguing against the regression-to-the-mean possibility. Human observers, in contrast, showed a different pattern. In a continuous recognition task (Bainbridge, 2017), degradation by both Gaussian noise and blur impaired recognition overall, consistent with prior work on memory for degraded images (e.g., Meng & Potter, 2008), but recognition of highly memorable images was resistant to degradation, whereas recognition of forgettable images declined. This effect was driven primarily by false alarms, reliably observed for noise and blur with low-memorability images, indicating that participants increasingly mistook novel low-memorability images for repeated ones. Global-matching models (Gillund & Shiffrin, 1984) predict that greater similarity to stored items raises false familiarity, so this finding prompts the development of a new model that accounts for item-to-item similarity.

Prior work has shown that noise-trained models mirror human behavioral and neural responses better than standard models (e.g., Jang et al., 2021; Jang & Tong, 2024; Thompson, 2021). We sought to test if conventional memorability models could similarly benefit from fine-tuning with noisy images. Our results suggest otherwise. Fine-tuning models on images randomly perturbed with Gaussian noise did not reliably improve the Spearman correlations with LaMem memorability scores, and for some models it was worse (e.g., ResMem and Vit_DINOv3). This is consistent with recent evidence that degradation training does not reliably produce human-like behavior even in object recognition. In a comparison of more than 200 DNNs, models trained on blurred or stylized images failed to match human recognition and were often less human-aligned than standard ImageNet-trained models (Kato & He, 2026). Why might noise training benefit classification in some studies but not memorability estimation? Research has shown that humans acquire noise robustness in object recognition through natural development (Bernstein et al., 2005) and through experimental training (Gold et al., 1999). Therefore, the model improvement in image classification under noise can be viewed as noise-training mimicking the extensive experiences human observers have in encountering stimuli under challenging visual environments. Importantly, recent developmental studies revealed that image memorability remains consistent across individuals at different stages of development (Almog et al., 2023), and that the memorability effect emerges as early as age four (Guo & Bainbridge, 2024). This suggests that the experience shaping memorability may be broad, early-life visual exposure rather than experience with degraded viewing specifically. Instead, the central question may be which image properties determine memorability and how noise alters them.

One property that may determine how noise shapes memorability is how similar images are to one another. It is well documented that memorization naturally relies on statistics of similarity and relational structure (e.g., Gillund & Shiffrin, 1984; McClelland & Chappell, 1998; Shiffrin & Steyvers, 1997), and the relational position and distinctiveness of an item determine how well it is remembered (Brown et al., 2007; Hunt & Lamb, 2001; Lukavsky & Dechterenko, 2017; Nosofsky, 1984). Therefore, models may need to account for the relational structure of the image set to match human memorability estimates and their tolerance to degradation. Indeed, recent work has challenged the notion of intrinsic memorability and instead argues that memorability is an emergent property of the relational structure and is dictated not by intrinsic pictorial features but rather by their distinctiveness in a given test set. Simpson and colleagues (2026) showed that model simulations on items with randomly generated features, and thus no intrinsic memorability, still produced consistent memorability differences driven by distinctiveness. In behavioral experiments, placing images among more or fewer similar items inverted their original memorability rankings, and changing which concepts were overrepresented in a subset of the THINGS dataset reversed which features predicted good memory. If memorability depends on an image’s position relative to other images, then noise, which alters image similarity, should change memorability in ways that feature-based conventional models miss.

Our results support this by developing a model that accounts for relational structure (GraphMem), as it was trained on graph-based image similarity rather than individual image features. Based on the Spearman correlation between model estimation and human data, GraphMem was the best model at predicting human recognition under noise, with the highest correlation with human performance in NoiseMem (ρ = .85 and .84). It was also the only model whose Slope Effect was not strongly negative (−.11 when trained on clean images) and the only one to turn positive (.57 when trained with noise) while every other model ranged from −.54 to −.82. This contrast highlights that the two measures capture different aspects of human-likeness. All models captured the overall memorability ranking under noise, but only GraphMem reproduced how noise changes memory across memorability levels. GraphMem trained with noise was the only model to match human data on both measures, and it was on the Slope Effect, the measure most diagnostic of how noise affects memory, that noise training produced its largest improvement. We propose that both outcomes stem from the information each model relies on when estimating memorability. Whereas conventional models estimate memorability from an image’s features alone, GraphMem estimates it from the image’s position in a similarity graph, which likely limits its accuracy in predicting memorability for individual images. Yet, noise degradation greatly affects image similarity, supported by typicality results, and thus its effect on memorability can be better characterized by GraphMem. The next natural step in developing models that better align with human performance is to consider both feature-based and relation-based information.

Are more memorable images processed more efficiently? Our overall results offer partial support to this account. Based on the efficient processing account of memorability (Bainbridge et al., 2026), memorability can be viewed analogously as signal quality, with more memorable images being stronger signals and thus processed faster and more efficiently. Accordingly, signal quality should be differentially affected by noise (i.e., stochastic resonance), and a similar pattern might be observed with memorability. Our human results are broadly consistent with this prediction. Although performance generally declined as noise increased, the magnitude of this decline varied with memorability, suggesting that memorability influences how effectively image information can be processed under degraded viewing conditions. However, this pattern differs from the stochastic-resonance-like effect observed in conventional DNNs. One possible explanation concerns the specific mechanism through which processing efficiency operates. Stochastic resonance occurs when noise can enhance the detectability of a weak, subthreshold signal by pushing it above a detection threshold (Simonotto et al., 1997). Whether memorability has a comparable threshold, and whether less memorable images fall below such a threshold, is unclear. Thus, the relationship between noise and memorability may not necessarily follow the same pattern as noise and perceptual detection. Furthermore, this divergence may reflect differences in how humans and models separate signals from noise. DNNs may be more susceptible to treating image-embedded noise as informative features and incorporating it into their representations alongside task-relevant signal, which could produce the stochastic resonance-like pattern. Human observers, in contrast, may have learned through extensive visual experience to filter out external noise and extract only the diagnostic information (i.e., the signal) (Dosher & Lu, 1998, 1999). Thus, the absence of stochastic resonance in humans does not necessarily contradict the efficient-processing account. Processing efficiency may instead be expressed as greater robustness to noise, with more memorable images retaining their processing advantage even as overall performance declines.

## Methods

### Stimuli

The image set was selected based on previous work on image memorability (Ma et al., 2024). Specifically, a set of images was chosen from the LaMem dataset (http://memorability.csail.mit.edu/index.html). This dataset contains 58,740 images from a variety of sources, each with a corresponding memorability score reflecting the probability of that image being recalled later. Twenty-eight images were randomly sampled from each of seven equally spaced memorability bins, or ranges of memorability scores (Δbin ≈ 0.10390; Bin 1, 1–0.89610; Bin 2, 0.89610–0.79220; Bin 3, 0.79220–0.68831; Bin 4, 0.68831–0.58441; Bin 5, 0.58441– 0.48051; Bin 6, 0.48051–0.37662; Bin 7, 0.37662–0.27273). Within each bin, scores were additionally constrained to be uniformly spaced. These images were then perturbed by five levels of zero-mean Gaussian noise, using the *imnoise* function in MATLAB (Gaussian σ = 0, 0.01, 0.05, 0.10, and 0.15; Fig. **5**), resulting in 980 images.

**Fig. 5.**
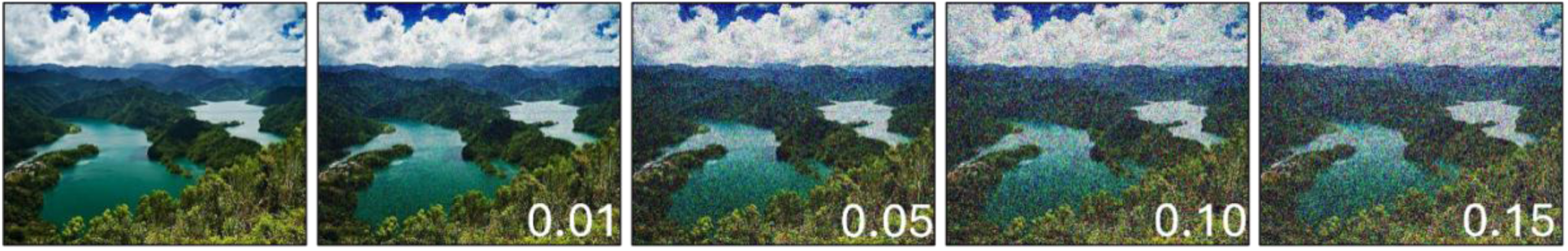
Example of image stimuli used. Numbers correspond to the sigmas of Gaussian white noise added to the image.

### Memorability models

We estimated memorability with three pretrained memorability models: ResMem (Needell & Bainbridge, 2022), a deep neural network trained to predict memorability on the LaMem dataset, based on a ResNet-150 architecture; ViTMem (Hagen & Espeseth, 2023), based on a Vision Transformer architecture; and AMNet (Fajtl et al., 2018), which uses a ResNet-50 architecture combined with three Long Short Term Memory (LSTM) steps. To test whether the human–model divergence generalized beyond these models, we additionally trained several state-of-the-art (SOTA) vision models, selected from the top models on the Hugging Face PyTorch Image Models leaderboard (https://huggingface.co/spaces/timm/leaderboard), on the LaMem dataset using standard methods. To do this, we used the test and train splits provided by the LaMem dataset. All models were built by taking the base ViTMem and swapping out the model backbone used for initializing the model. For all testing, we used pretrained models that had been evaluated on ImageNet 1k, which were then fine-tuned using LaMem. All models were trained in PyTorch using the Adam optimizer and mean squared error (MSE) for the loss function. For the final SOTA models selected, we specifically chose (“convnext_tiny.fb_in22k”, “eva02_tiny_patch14_336”, “vit_base_patch16_clip_224”, “vit_small_patch16_dinov3.lvd1689m”, and “vit_base_patch16_224.mae”). For training with noise, each image in the LaMem dataset was randomly perturbed with zero mean Gaussian noise with σ chosen from a uniform distribution between 0 and 0.15. For retraining ViTMem, ResMem, and AMNet, a similar approach was taken by retraining each model on the LaMem dataset with noise perturbations.

### Attention-map and typicality analysis

AMNet combines an ImageNet-pretrained ResNet-50 feature extractor with a soft attention mechanism and an LSTM that, over three recurrent steps, produces a spatial attention map at each step (Fajtl et al., 2018). These attention maps indicate which image regions the model emphasizes when estimating memorability. We extracted these maps from the final (third) LSTM step for each image in its original form and at each noise level and calculated the entropy of each attention map to represent its degree of dispersion.

For typicality, we used the ImageFluency toolbox for R (Mayer, 2026). Following Mayer and Landwehr (2018), typicality was operationalized as the pixel-wise correlation between a given image and the average representation of the image set (i.e., the pixel-wise mean of all images in the set). Scores can range from −1 (inversely typical) through 0 (not typical) to 1 (perfectly typical), with higher values indicating that an image more closely resembles the prototype of the set; for our analysis, we used the absolute value of typicality. Typicality scores were calculated separately for each noise level.

### GraphMem

We trained our GraphMem model on the LaMem dataset, using the split 1 test provided (Khosla et al., 2015). Each image in the training set was resized to 224x224 pixels and passed through a ViT-B/16 model, using the same base model as in ViTMem (vit_base_patch16_224) with default weights and pretrained on ImageNet-21k and fine-tuned on ImageNet-1k. The classification head was removed, and the 768-vector embeddings from the final layer were extracted. These embeddings were then used to construct a k-nearest-neighbor graph using the Python *scikit-learn* library, using the cosine similarity of each pair of embeddings to find the k=10 images with the highest similarity to each image. This graph was then fed into a two-layer Graph Convolutional Network (GCN; van dan Berg et al., 2017) as implemented in the PyTorch Geometric library (version 2.8.0; Fey & Lenssen, 2019). The two-layers in this network correspond to how each image is linked in two steps to the rest of the nearest neighbors graph; that is, for each image, the GCN learns its similarity to the k=10 nearest neighbors at the first layer, and then how each of those images relate to *their* k=10 nearest neighbors at the second layer. Each layer was followed by a ReLU function and had 256 units. These were then fed into a linear output unit to produce a memorability score for each image.

For training, the GCN used the AdamW optimizer with MSE as the loss function across the entire batch for 500 epochs. Every 10 epochs, the validation set from LaMem split 1 was added to the graph with nearest-neighbor link recomputed for the combined set. The Spearman correlation was then calculated between actual and estimated memorability scores. Across the entire 500 epochs, we retained the weights with the highest correlation, which was then applied to the test set (reported in Table 1). For both training and testing, we ensured that the 196 images from NoiseMem were not included in the split 1 set. Next, the 980 images from NoiseMem were tested by again applying them through the same pipeline as for the validation and test sets; a similar procedure was also used for predicting memorability scores for BlurMem. We additionally, observed that splitting the 980 images up into separate sets, or running them individually, produced similar results.

For training the GCN with noise, we encountered a particular challenge, in that adding noise changed the training graph structure, depending on which images received which amounts of noise; for example, if high memorability images randomly received more noise than low memorability images, this would produce a different graph structure than if the opposite occurred. While the large size of the training set likely led to noise effects averaging out across memorability levels, we sought to control and measure these effects. To do this, we trained the model in 100 separate instances, varying the random seed used for both adding noise and for the starting weights of the model. This was done to mimic the 100 subjects collected for both the NoiseMem and BlurMem human datasets (Rafiei et al., 2024). We additionally ensured that noise was uniformly distributed across memorability by adding noise in batches to each memorability decile. The memorability predictions were then averaged across the 100 training sets for the 980 images in the NoiseMem and BlurMem sets.

### Human experiments

#### Participants

##### NoiseMem experiment

A total of 100 participants were recruited for an online experiment (mean age = 20.3 years, SD = 1.92; 57.5% female, 25.6% male, 16.9% other). All participants were drawn from the undergraduate pool of George Mason University and took part for course credit, with all protocols approved by the Institutional Review Board. All participants provided informed consent and had normal or corrected-to-normal vision. Twelve participants did not provide any responses throughout the experiment and were excluded from the following analyses. The experiments were programmed using PsychoPy (version 2020 and above), and the online experiments were conducted using the Pavlovia platform (www.pavlovia.org).

##### BlurMem experiment

A total of 101 participants were recruited for an online experiment (mean age = 22.6 years, SD = 6.1, 74.3 % female, 23.8% male, 2% other). All participants were drawn from the undergraduate pool of George Mason University and took part for course credit, with all protocols approved by the Institutional Review Board. All participants provided informed consent and had normal or corrected-to-normal vision, and none took part in the NoiseMem experiment. Ten participants did not provide any responses throughout the experiment and were excluded from the following analyses.

#### Stimulus presentation

We used the same 196 LaMem images described above, 28 images sampled from each of seven equally spaced memorability bins, together with their five noise-perturbed versions (0, 0.01, 0.05, 0.10, and 0.15), yielding a pool of 980 image files. Ground-truth memorability scores were taken from the LaMem database and were not re-estimated for the present experiment, so the bin assignments used in the human analyses are identical to those used in the model analyses. Because each participant viewed a given image at only one noise level, these 980 images served as a shared pool from which each participant’s individual stimulus set was drawn. Images were presented at 400x400 pixels against a gray background.

#### Procedure

##### NoiseMem experiment

Participants completed a modified version of the “memory game” continuous recognition task (Khosla et al., 2015; **Fig. 2a**), in which images were presented sequentially (750 ms) followed by a 1 s gap, during which participants indicated via key press whether the image just shown was a repeat of one they had seen earlier in the sequence. A total of 430 images were viewed during the experiment. Each participant saw 240 target presentations (120 unique images, each shown once at encoding and once as a test repeat 35–139 images later) and 190 filler presentations. Most filler images (114) were presented only once, providing spacing between a target and its repeat, whereas 38 fillers were presented twice, with 0 to 3 intervening images between their two presentations; these vigilance repeats served as attention checks. In addition, every image (both targets and fillers) was presented at one of five levels of visual noise, with a target’s noise level held constant across its study and test presentations, allowing recognition memory to be assessed as a function of image degradation. Each participant only viewed a given image at a single noise level, and a counterbalancing algorithm (Goetschalckx et al., 2019; https://github.com/LoreGoetschalckx/MemoryGame) was used to ensure that each participant received a unique sequence of targets and fillers in randomized order, drawn from a shared item pool so that images received balanced exposure across both item role and noise level across the entire sample.

##### BlurMem experiment

We used the same set of 196 images but now perturbed with five levels of visual blur, using the MATLAB function *imgaussfilt* (Gaussian σ 0–10; **Fig. 4a**). Participants completed the continuous recognition “memory game,” identical to the NoiseMem experiment.

### Statistical analysis

All analyses followed the same general two-step procedure. First, for each dependent variable, we quantified its change across perturbation levels by fitting a simple linear model across the five noise (or blur) levels and extracting the slope. For model analyses, slopes were obtained for each image (estimated memorability, attention-map entropy, or typicality). For human analyses, slopes were obtained for each participant on hit rate (HR), false alarm rate (FA), and corrected recognition (CR), where CR was calculated by subtracting the false alarm rate from the hit rate for target images. Second, these slopes were submitted to a one-way ANOVA with memorability bin as the factor to determine whether the change in estimated memorability or human performance across perturbation levels varied as a function of memorability. Memorability bin was treated as a between-image factor for model analyses and as a within-participant factor for human analyses (repeated-measures ANOVA). To determine the direction of these effects, we additionally regressed the slopes on memorability bin, coded as a continuous variable. For human data, we used a linear mixed-effects model with memorability bin as a fixed effect and participant as a random intercept; for model outputs, we used a simple linear regression with memorability bin as the predictor. The sign and significance of the bin coefficient indicate whether perturbation slopes increase or decrease as memorability bin increases (i.e., as images become less memorable).

## Data Availability

All behavioral data and stimuli are publicly available on the Open Science Framework at https://doi.org/10.17605/OSF.IO/W5JTM. The GraphMem model is available on GitHub at https://github.com/wimartin/GraphMem.

